# A role for the Tgf-β/Bmp co-receptor Endoglin in the molecular oscillator that regulates the hair follicle cycle

**DOI:** 10.1101/230771

**Authors:** María I. Calvo-Sánchez, Elisa Carrasco, Sandra Fernández-Martos, Gema Moreno, Carmelo Bernabeu, Miguel Quintanilla, Jesús Espada

**Author notes:** Address for correspondence: Jesús Espada, Experimental Dermatology and Skin Biology Group. Ramón y Cajal Institute for Biomedical Research (IRYCIS), Ramón y Cajal University Hospital, Colmenar Viejo Rd. Km 9,100, 28034 Madrid, Spain.

## Abstract

The hair follicle is a biological oscillator that alternates growth, regression and rest phases driven by the sequential activation of the proliferation/differentiation programs of resident stem cell populations. The activation of hair follicle stem cell niches and subsequent entry into the growing phase is mainly regulated by Wnt/β-catenin signalling, while regression and resting phases are mainly regulated by Tgf-β/Bmp/Smad activity. A major question still unresolved is the nature of the molecular switch that dictates the coordinated transition between both signalling pathways. Here we have focused on the role of Endoglin (*Eng*), a key coreceptor for members of the Tgf-β/Bmp family of growth factors.

Using an *Eng* haploinsufficient mouse model we report that *Eng* is required to maintain a correct follicle cycling pattern and for an adequate stimulation of hair follicle stem cell niches. We further report that β-catenin binds to the *Eng* promoter depending on Bmp signalling. Moreover, we show that β-catenin interacts with Smad4 in a Bmp/Eng dependent context and both proteins act synergistically to activate *Eng* promoter transcription. These observations point to the existence of a growth/rest switching mechanism in the hair follicle that is based on an Eng-dependent feedback crosstalk between Wnt/β-catenin and Bmp/Smad signals.

## INTRODUCTION

The hair follicle can be considered as a complex mini-organ with a distinctive morpho/functional identity that continuously cycles through growing (anagen), regression (catagen) and resting (telogen) phases (Fuchs, 2007; Hsu et al., 2014; Mu et al., 2001; Plikus et al., 2008; Schneider et al., 2009). The activity of the whole hair follicle is controlled by specialized populations of stem cells mainly located in the bulge region, site of insertion of the arrector pili muscle, and in a morphologically distinguishable structure below the bulge known as the hair germ (Cotsarelis et al., 1990; Greco et al., 2009; Hsu et al., 2014; Morris et al., 2004; Rompolas et al., 2013; Sun et al., 1991; Tumbar, 2004). Hair follicle stem cells periodically leave their quiescent state and are induced to proliferate, expand and differentiate in response to oscillating signals originated in the surrounding niche (Fuchs, 2007; Hsu et al., 2014; Plikus et al., 2008). At the onset of the anagen phase, hair germ stem cells are the first to proliferate giving rise to a transit amplifying population of progenitor cells that migrate downwards ultimately forming the hair matrix region. Matrix cells can in turn proliferate and differentiate upwards giving rise to the hair shaft and the surrounding inner root sheath. Bulge stem cells give raise to the outer root sheath, a cell population retaining stemness characteristics that envelops the matrix transit amplifying cell population at the differentiating core of the hair follicle (Hsu et al., 2014). Once matrix cells stop their proliferation and differentiation programs, hair follicle enters in the catagen phase and its lower part undergoes a rapid involution triggered by the activation of matrix cell apoptosis. Eventually, the lower part of the hair follicle is reduced to a single epithelial strand, bringing the dermal papilla into close proximity to the bulge region (Schneider et al., 2009). This process is followed by the entry of the hair follicle in the resting or telogen phase, where dermal papilla cells become progressively competent to activate again the bulge region in response to specific signals. The cyclic nature of hair follicle activity, and the fact that this activity depends on a well-defined population of stem cells, makes this structure a suitable biological model to investigate new modes of functional regulation of adult stem cell dynamic micro-environments in mammals.

Different paracrine signalling pathways have been directly involved in the regulation of the hair follicle growth cycle (Plikus et al., 2008). Among them, Wnt/β-catenin and Bmp/Smad are the best characterized to date (Supplementary Fig. 1). Wnt/β-catenin signalling regulates the onset and progression of the anagen (Schneider et al., 2009; Shimomura and Christiano, 2010), while activation of Bmp/Smad signalling during mid anagen is involved in the entrance of the hair follicle in the catagen and telogen states (Oshimori and Fuchs, 2012; Plikus and Chuong, 2008; Plikus et al., 2008). The activation of the Bmp/Smad pathway is currently used to determine the hair follicle stages, using the expression levels of Bmp2 and Bmp4 as late anagen and refractory telogen markers (Plikus et al., 2008). Progressive activation of Bmp/Smad signalling in an out-of-phase pattern with respect to Wnt/β-catenin signalling further divides the telogen into two sub-phases, refractory (expression of Bmp) and competent (absence of Bmp) (Plikus et al., 2008) (Supplementary Fig. 1). During the refractory telogen, hair follicles are unable to respond to anagen re-entry stimulation even in the presence of Wnt/β-catenin signalling. Throughout the second part of the telogen phase, attenuated Bmp signalling allows the anagen re-entry wave to propagate, and hair follicles are competent to respond to Wnt/β-catenin signalling. The whole system can be accurately described as a biological oscillator regulated by negative feed-back mechanisms (Sasai, 2013) in which two broadly defined interacting cell populations, bulge stem cells and dermal papilla niche cells, alternate active and inactive states depending on a basic stimulatory signal (Wnt/β-catenin) that is cyclically regulated by a negative signal (Bmp/Smad). On the other hand, taking into account that the activation of the Wnt pathway strengthens Bmp signalling (Baik et al., 2016), a major unanswered question is the nature of the molecular mechanisms that so precisely regulate the coordinated switch between growth and rest, and backwards, in the hair follicle stem cell niche.

Here we have focused on the role of Endoglin (Eng), an essential co-receptor of Bmp growth factors, in the regulation of the hair follicle cycle. In the mouse, *Eng* is expressed early during embryonic development on mesenchymal tissue derived from the endocardium and also in the vascular endothelium, playing a critical role in cardiovascular system development and homeostasis (Bourdeau et al., 1999). After birth, *Eng* is expressed mainly in endothelial cells, and, to a lesser degree, in macrophages, fibroblasts, vasculature muscle cells, mesenchymal and haematopoietic stem cells, blood cells, and also in several regions of the skin, such as the interfollicular epithelium and hair follicles (Quintanilla et al., 2003; Supplementary Fig. 2). Mutations in *Eng* gene are associated to the hereditary haemorrhagic telangiectasia vascular dysplasia, termed HHT1 (Kapur et al., 2013; López-Novoa and Bernabeu, 2010; McAllister et al., 1994). Eng is also involved in skin regeneration during wound healing (Pérez-Gómez et al., 2014) and can supress keratinocyte proliferation in early stages of a multistage mouse skin carcinogenesis model, driving malignant progression, invasion and metastasis, in later phases (Pérez-Gómez et al., 2007; Quintanilla et al., 2003). These observations point to important roles for *Eng* in the regulation of skin stem cell niches and in the maintenance of skin homeostasis similar to the role of this protein in the hematopoietic system (Baik et al., 2016).

## METHODS

### Cell culture procedures

The feeder-independent E14 mouse embryonic stem (ES) and human embryonic kidney 293T cell lines were used in this study. E14 cells (a gift from T. Rodriguez, Imperial College, London) were grown on 0,1% gelatin-coated flasks in GMEM supplemented with 10% FCS, NEAA, L-Glutamine, pyruvate, β-mercaptoethanol (all from Gibco) and 1,000 U/ml LIF (Millipore), and splitted 1:8 or 1:10 every 2–3 days using Trypsin-EDTA (Invitrogen) (Cambray et al., 2012; Smith, 1991). 293T cells were cultured in DMEM supplemented with 10% FCS, L-Glutamine and antibiotics, and divided 1:10 or 1:20 every 2-3 days using Trypsin-EDTA (Invitrogen). Both cell lines were maintained at 37°C in a 5% CO_2_ humidified atmosphere.

For *Eng* RNA interference in E14 cells, a siRNA cocktail containing 4 different oligonucleotides (50 nM each) targeting different regions of the coding sequence was used (Qiagen, S1009938-11, -18, -25, -32). Inhibition of *Eng* expression was evaluated by qRT-PCR. For protein over expression experiments, pcDNA3-Smad4 (Kang et al., 2005) and pCI-neo-mutant β-catenin (S33Y) (Espada et al., 2005; Morin et al., 1997) constructs were transiently transfected in 293T cells. Protein expression levels were evaluated by immunoblotting. For *Eng* promoter expression assays, pCD105 (−50/+350) and pCD105 (−2450/+350) reporter constructs derived from the human *ENG* promoter were used as described (Botella et al., 2002). *Luciferase* reporter activity was determined in whole cell lysates using the dual GLO-luciferase assay kit (Promega) and *Renilla* expression vector as an internal control, following the manufacturer's instructions. The empty luciferase vector pXP2 was used as control. The amount of DNA in each transfection was normalized using the corresponding empty vector. In all cases, transfection assays were performed using Lipofectamine^®^ 2000 transfection reagent (Life Technologies) following manufacturers' instructions.

### Animal Procedures

Generation, maintenance and genotyping of an *Eng*^+/−^ mouse strain on a C57Bl/6 background have been previously described (Bourdeau et al., 1999). Animals were kept in ventilated rooms under lighting (12-h light, 12-h dark cycle) and temperature controlled conditions, and allowed feed and water *ad libitum*. All experiments were conducted in parallel in *Eng*^+/−^ and *Eng*^+/+^ littermate mice aged 1-5 months. All experimental procedures were conducted in compliance with 2010/63/UE European guidelines.

Induction of hair growth was basically performed at two different points during the hair follicle cycle: anagen/refractory telogen transition (postnatal day 50) and competent telogen/anagen transition (postnatal day 90) (Plikus and Chuong, 2008; Plikus et al., 2009). Haired back skin regions of at least 3 mice of each genotype and each hair cycle phase were clipped as described (Carrasco et al., 2015). Progression of hair growth was sequentially monitored and imaged daily until most animals of one genotype completed fully hair coating. Hair clipping was chosen over plucking or shaving to avoid wounding that can potentially interfere with normal hair growth (Chase, 1954; Plikus and Chuong, 2008).

Activation of skin stem cell and transit amplifying cell proliferation and mobilization in back and tail skin was performed by sequential (3 times) topical application of 3 doses of 20 nM 12-O-Tetradecanoylphorbol-13-acetate (TPA, Sigma-Aldrich) (Braun et al., 2003; Espada et al., 2008; Fürstenberger and Marks, 1980). Skin stem cell proliferation was also activated by induction of a transient production of reactive oxygen species (ROS) in the tissue as previously described (Carrasco et al., 2015; Fischer et al., 1986).

### Immunological procedures

Antibodies used in this work are listed in Supplementary Table I.

For immunolocalization of proteins in histological sections, pieces of dorsal skin were fixed in pH 7.0 buffered 3.7% formaldehyde and processed for histology. Histological sections were stained with the indicated antibodies after permeabilization in 0.1% Triton X-100, except in the case of Endoglin detection, and blocked in 0.5% BSA. The immunofluorescent signal was revealed using HRP-coupled secondary antibodies and TSA Plus Cyanine 3 (Perkin Elmer) for signal amplification, following the manufacturer's instructions. Histological sections were also stained with standard haematoxylin-eosin for routine evaluation of tissue morphology.

For the preparation of tail epidermis whole-mounts, tails were clipped and the skin was peeled and treated with 5 mM EDTA. Intact sheets of epidermis were separated from the dermis, and fixed in pH 7.0 buffered 3.7% formaldehyde. For BrdU detection in whole-mounts, intact sheets of epidermis were washed in PBS several times in order to remove excess of formaldehyde, and treated with HCl 5N for nuclear acid hydrolysis, followed by TBE (Tris-Borate-EDTA) for neutralization. After blocking and permeabilization with PBT buffer (0.5% Triton X-100 and 0.2% gelatine in PBS), epidermis sheets were stained with FITC-conjugated mouse monoclonal anti BrdU, as previously described (Braun et al., 2003). For apoptosis detection in hair follicles of tail epidermis, TUNEL Label (Roche) assay was performed.

Confocal images were obtained in Leica TCS SP2 and SP5 AOBS spectral confocal microscope and processed using the FIJI software (Image J 1.49 National Institutes of Health, Bethesda, MD).

For immunoblotting, cell pellets or whole skin samples were homogenized in RIPA (25 mM Tris-HCl pH 7.6, 150 mM NaCl, 1% NP-40, 0.1% SDS) or SDS buffer (50 mM Tris-HCl pH 6.8, 2% SDS, 10% glycerol, 1% β-mercaptoethanol, 12.5 mM EDTA) containing protease and phosphatase inhibitors (2 μg/ml aprotinin, 2 mM PMSF, 2 μg/ml leupeptin and 2 mM sodium orthovanadate, 2 mM ß-glycerophosphate, 5 mM NaF, all from Sigma-Aldrich). Skin samples were fully disaggregated using scissors and a Polytron^®^ homogenizer (PT 1200 E, Kinematica), and the same amount of protein of each mouse was loaded in Laemmli buffer. Proteins were resolved in a 7.5% SDS-PAGE system, and were transferred to a PVDF membrane, which was blocked and stained with the indicated primary and secondary HRP conjugated antibodies. Finally, HRP activity was detected using the ECL-chemiluminescent kit (Amersham) in accordance with the manufacturer's instructions.

For Chromatin Immunoprecipitation (ChIP) assays the ENCODE and modENCODE guideline standards for ChIP experiments and data analysis (ENCODE Consortium V 2.0, 2011) were followed. Cell pellets and skin samples disaggregated with scissors were fixed in 1% formaldehyde-PBS, disaggregated and processed in ChIP lysis buffer (1% SDS, 10 mM EDTA,50 mM Tris HCl pH 8.0) with protease an phosphatase inhibitors (2 μg/ml aprotinin, 2 mM PMSF, 2 μg/ml leupeptin and 2 mM sodium orthovanadate, 2 mM β-glycerophosphate, 5 mM NaF, all from Sigma-Aldrich). Supernatants were sonicated and cellular debris removed by centrifugation. To evaluate proteins binding to the *Eng* promoter, supernatants were incubated with the indicated primary antibodies, DNA-protein interaction was reversed and qRT-PCR or conventional PCR were performed using the DNA *mEng* primers Fwd3 and Rvs3 (Supplementary Table II).

For protein co-immunoprecipitation assays, cell pellets or skin samples were homogenized in IPH buffer (50 mM Tris-HCl pH 7.4, 100 mM NaCl, 10% glycerol, 5mM EDTA, 0.15% Triton ×100) containing protease and phosphatase inhibitors as described before. Skin samples were disaggregated using scissors, and the supernatant was incubated O/N with the indicated antibodies and the equivalent IgG as a negative control. Proteins were resolved in a 7.5% SDS-PAGE system and transferred to PVDF membranes that were incubated with the indicated primary and secondary antibodies.

### Gene expression procedures

For RNA extraction, E14 cell cultures coming from the treatments described above, or dorsal skin of at least three mice at each time point and genotype, RNeasy mini kit and RNase-Free DNase Set (both form Qiagen) were used. Skin tissue was homogenized using TriPure™ isolation Reagent (Roche), disaggregated and processed using scissors and a Polytron^®^ homogenizer (PT 1200 E, Kinematica). For reverse transcription, MLV enzyme (Promega) was used, loading the same amount of RNA. qRT-PCR assays, or semiquantitative PCR, were performed for gene expression analysis, using Power SYBR Green (Applied Biosystems), or REDExtract-N-Amp™ PCR ReadyMix™ (Sigma Aldrich), respectively, following manufacturer's instructions. Specific primers, detailed in Supplementary Table II, were designed among different exons, thereby avoiding residual genomic DNA amplification, for semiquantitative or quantitative transcript detection.

Microarray experiments were performed using Mouse Gene Expression 4×44K Microarray Kit G4122F (Agilent technologies, Wilmington, DE). RNA was isolated using RNAesy Extraction Kit (QIAGen, Germany). RNA was labeled and array hybridized using the Low RNA Linear Amplification Kit and the In Situ Hybridization Kit Plus (Agilent technologies, Wilmington, DE) respectively. After hybridization and washing, the slides were scanned in an Axon GenePix Scanner (Axon Instruments Inc., Union City, CA) and analyzed using Feature Extraction Software 10.1 (Agilent technologies, Wilmington, DE). Two different RNA samples obtained from each *Eng* modified cell line were labeled with Cy5-dUTP. The RNA samples extracted from wild type cells were marked with Cy3-dUTP (Amersham, Sweden). Two additional hybridizations were performed using the reciprocal fluorochrome labeling. The genes whose expression was up or downregulated at least 2fold in *Eng*^+/−^ with respect to control cells were selected for analysis. Microarray raw data tables have been deposited in the Gene Expression Omnibus under the accession number of GSEXXX (submitter G. M.-B.).

### Statistical analysis

Quantifications of LRC in the bulge region were performed on confocal images (30 hair follicles/animal, 3 animals/group). Comparisons between groups were performed by Student's t-test using the SPSS 15.0 software (IBM, Spain). For statistical analyses of gene expression data, an unpaired t-test was applied, setting P≤0.05 as limit for significance. Quantitative realtime reverse-transcriptase–PCR data were analyzed using a comparative CT method, by using 18S ribosomal RNA expression as an internal control. Gene expression changes were represented as 2^−ΔCt^ values of *Eng*^+/−^ and control groups mean values at different time points during the second postnatal hair growth cycle. For quantification of hair regeneration, day-to-day digital images were analyzed. The bold area was quantified using the FIJI software. The area under the curve was calculated separately for each mouse, and means were compared by the Student's t-test. For statistical analysis of *Luciferase* reporter activity between the experimental samples, an unpaired t-test was applied setting P≤0.05 as limit for significance. Transactivation assay results were expressed as the ratio between Luciferase activity and *Renilla* expression vector as an internal control.

## RESULTS

### Eng shows a hair follicle cycle-dependent expression pattern in mouse skin that is deregulated in *Eng* haploinsufficient mice

We first sought to determine the expression pattern of *Eng* during the hair follicle cycle in wild-type (*Eng*^+/+^) C57Bl/6 mice, using as experimental framework the second of the two coordinated hair follicle cycles that take place in this biological model. This second cycle, ranging from 30 to 100 postnatal days, is the more lengthy and easily manipulable from an experimental perspective We found that the *Eng* mRNA exhibited a hair cycle-dependent expression pattern in *Eng*^+/+^ mice, showing a very low expression level during the anagen phase, a gradual increase, starting at the onset of the telogen (postnatal day 50, anagen/refractory telogen transition), to reach a maximum peak at the competent telogen/propagant anagen transition (postnatal day 90), followed by a drastic decrease henceforth (Fig. 1A). This result was broadly confirmed by the analysis of the Eng protein expression pattern in the skin (Fig. 1B). Interestingly, such expression pattern perfectly fits with the profile of master feedback target regulators of the hair follicle cycle predicted by a robust mathematical model that describes hair follicle dynamics as the result of coupled mesenchymal and epithelial oscillators, and that, in fact, identifies Eng as one of those potential targets (Tasseff et al., 2014).

**Figure 1.**
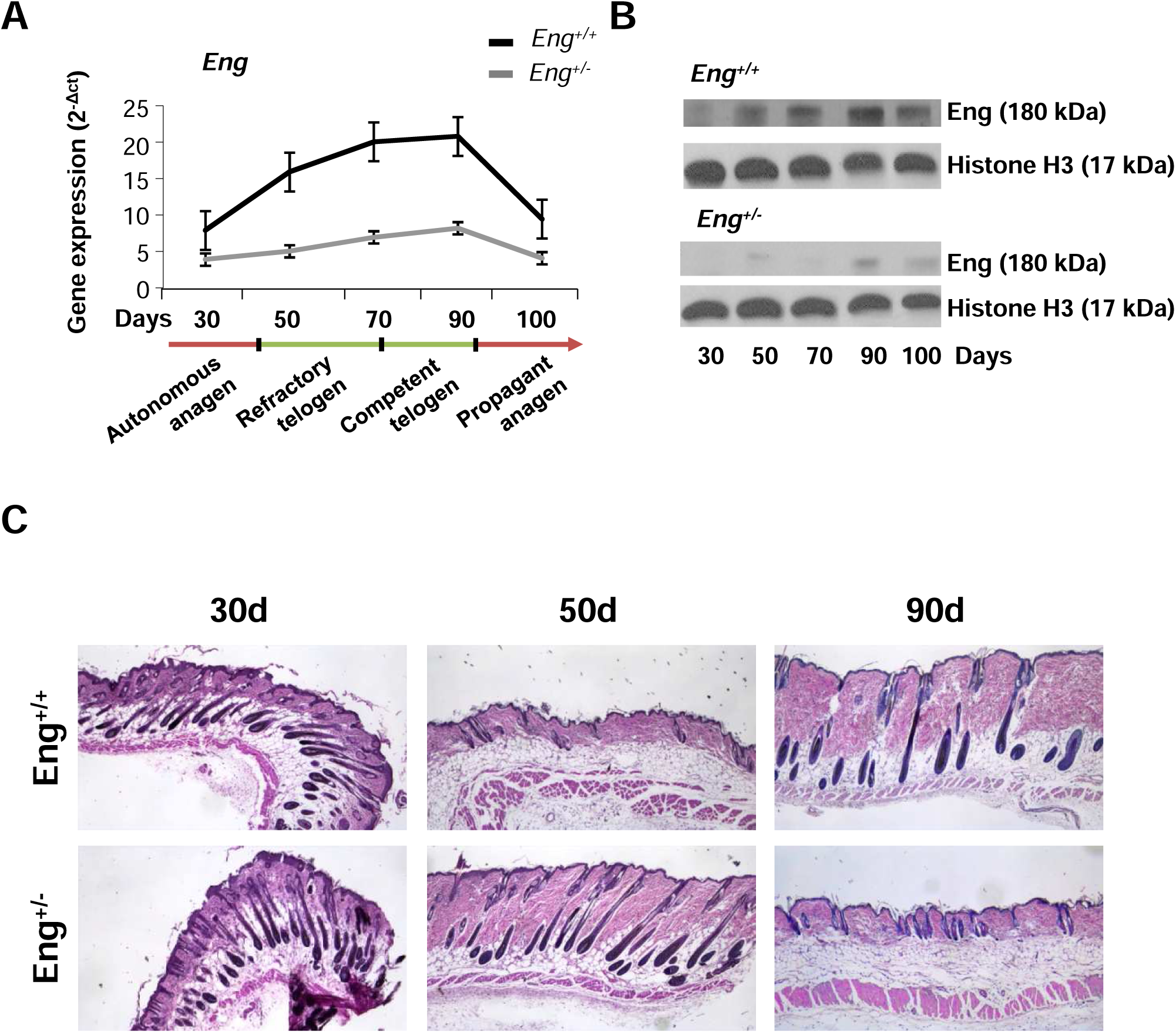
The *Eng* cyclic expression pattern in mouse skin is deregulated under *Eng* haploinsufficienciecy resulting in a delayed entry into the refractory telogen phase. **A)** *Eng* mRNA expression quantification by qRT-PCR, normalized to 18S rRNA, in *Eng*^+/+^ and *Eng*^+/−^ mouse dorsal skin at different time points (postnatal days) of the hair growth cycle, showing the hair follicle growth phase-dependent cyclic expression pattern of this gene. The mean ± SE was represented (n=3 in each time point). **B)** Immunoblot analysis of Eng protein expression, and histone H3 as a loading control, in *Eng*^+/+^ and *Eng*^+/−^ mouse dorsal skin at the indicated time points (postnatal days) during the hair follicle cycle. **C)** Morphology of *Eng*^+/+^ and *Eng*^+/−^mouse dorsal skin at the indicated time points (postnatal days), showing hair follicles in full-length vertical orientation in histological sections stained with haematoxylin-eosin. A significantly delayed entry into the refractory telogen (postnatal day 50) is observed in *Eng*^+/−^ animals. Black bars represent average hair follicle length in each time point. Scale bar: 200 μm.

These observations prompted us to investigate the effect of a functional decrease of Eng in the skin. To this end we used C57Bl/6 mice lacking a copy of the *Eng* gene (*Eng*^+/−^). Haploinsufficient *Eng*+^/−^ mice in a C57Bl/6 background are essentially equivalent to wild-type animals with respect to physiopathology, behaviour, fertility and life expectancy. Clinical signs of HHT are almost absent in these animals (Bourdeau et al., 1999; Quintanilla et al., 2003). Moreover, the *Eng*^+/−^model also ensured the effective and functional reduction of Eng in the tissue. As expected, *Eng*^+/−^ mice showed a drastic reduction of Eng expression in the skin and a concomitant loss of a cyclic patterning during the hair follicle cycle (Fig. 1A, B). Although no differences were observed in hair follicle formation and dynamics during early postnatal development, a striking delay in the onset of the second telogen phase (postnatal day 50 in wild-type animals) was observed in *Eng*^+/−^ mice as compared to control *Eng*^+/+^ littermates (Fig. 1C). The altered pattern was maintained through all the second telogen phase so that Eng^+/−^ mice entered in the subsequent anagen phase by day 90 while *Eng*^+/+^ littermates were still in telogen (Fig. 1C). Such unusually modified hair follicle cycling was consistently observed in 97% of analysed animals (n=50). This result suggests that Eng plays a central role in the regulation of telogen entry, as predicted for feedback target regulators in the coupled dual oscillator model of hair follicle dynamics (Tasseff et al., 2014).

### The molecular signals that dictate the anagen-telogen transition are out-of-phase in *Eng* haploinsufficient mice

As Eng is a co-receptor of the Bmp cytokine family and, particularly Bmp4 is directly involved in the regulation of telogen entry, being used as a marker of late anagen and refractory telogen (Plikus et al., 2008), we investigated the expression pattern of this gene during the hair follicle cycle in *Eng*^+/−^ mice as compared to control *Eng*^+/+^ littermates. As expected, in wild-type animals Bmp4 showed high expression levels in the late anagen/refractory telogen transition that gradually decreased with progression through the hair follicle cycle and this pattern was not significantly altered in haploinsufficient conditions (Fig. 2A) indicating that the background pattern of molecular inputs that define the entry/exit of the hair follicle into different growth phases is not altered in *Eng* haploinsufficient conditions. We further analysed the expression pattern of *Id1*, a transcriptional target of Bmp4 signalling in the skin (Ahmed et al., 2011). Interestingly, we found that *Id1* showed an expression pattern resembling the profile of *Bmp4* in *Eng*^+/+^ animals, but this pattern was completely abrogated in *Eng*^+/−^ littermates (Fig. 2A). These results indicate that Eng is directly involved in the transmission of Bmpsignals in the skin.

**Figure 2.**
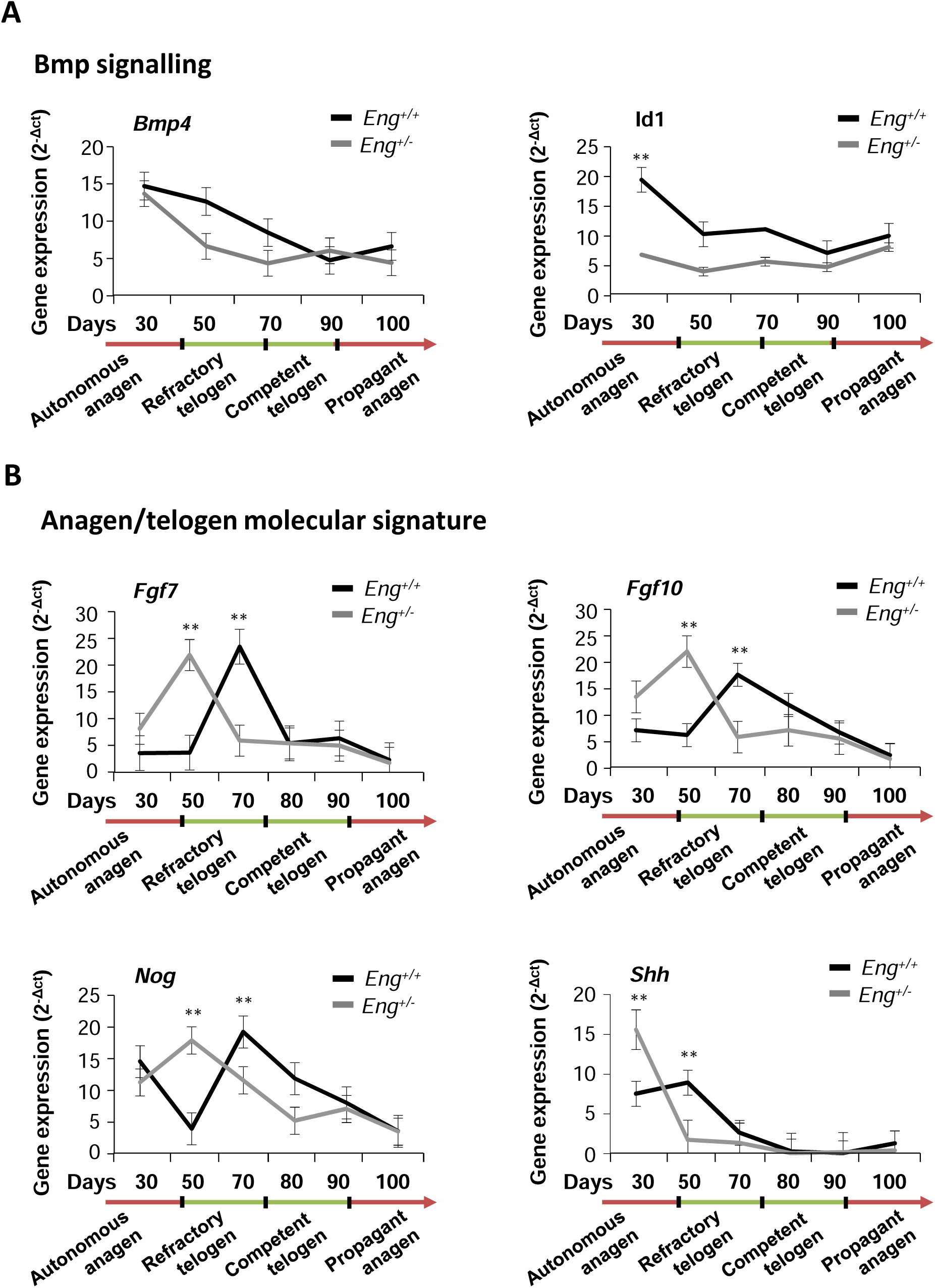
Disruption of Bmp signalling in the skin is associated with a premature out-of-phase expression of factors implicated in the regulation of hair follicle phase transitions in *Eng* haploinsufficient mice. **A)** *Bmp4* and *Id1* mRNA expression in *Eng*^+/+^ and *Eng*^+/−^ mouse dorsal skin at different points of the hair growth cycle, showing the loss of cyclic expression pattern of *Id1*, but not *Bmp4* in *Eng*^+/−^ animals. **B)** *Fgf7*, *Fgf10*, *Noggin* and *Shh* mRNA expression in *Eng*^+/+^ and *Eng*^+/−^ mouse dorsal skin at different points of the hair growth cycle, showing the premature out-of-phase expression of all these factors in *Eng*^+/−^ animals during autonomous anagen/refractory telogen transition. In all cases, gene expression was quantified by qRT-PCR, normalized to 18S rRNA and the mean ± SE was represented (n≥3 in each time point).

Next we wondered if the defective transmission of Bmp signals in *Eng*^+/−^ animals may disturb the molecular framework that modulates anagen/telogen/angen transitions. To this end, we investigated the gene expression patterns of key factors required to induce the re-entry of the hair follicle into the growing phase, including Fgf7, Fgf10, Nogging and Shh (Hsu et al. 2014 and references therein). Notably, we found that all these factors recurrently showed a premature out-of-phase expression patern in *Eng*^+/−^ mice as compared to control *Eng*^+/+^ littermates (Fig. 2B). This result is consistent with the deregulated hair follicle cycling pattern observed in these animals (Fig 1C) and suggest that Eng haploinssufiency impairs the molecular oscillator that dicates the alternation of Wnt/β-catenin and Bmp/Tgf-β/Smad signalling that occurs during the hair follicle cycle.

### Eng is required for correct telogen entry/exit timing during the hair follicle cycle

Our results suggest that Eng plays important roles in the anagen/refractory telogen and competent telogen/anagen transitions during the hair growth cycle. In this context, we hypothesized that a physiological stimulus to activate hair growth in the anagen/refractory telogen transition (postnatal day 50), when the hair follicle is unable to respond in normal conditions to stimulatory signals, or in the competent telogen/anagen transition (postnatal day 90), when the hair follicle is now ready to respond to growth signals (Plikus et al., 2008; Plikus et al., 2009), should have different responses in *Eng*^+/+^ and *Eng*^+/−^ littermates.

To test this hypothesis we performed a series of hair clipping experiments in the back skin of sample animals at 50 or 90 days after birth. As expected by the high expression of Bmp4, clipping stimulation in wild-type animals resulted in a delayed response of hair follicle growth during the anagen/refractory telogen transition, as compared to the rapid induction observed during the competent telogen/anagen transition (Fig. 3A), coinciding with low expression of Bmp4 and according to the molecular background of the effectors in the tissue in each time point. Thus, these animals showed a strong activation of Bmp/Smad signalling and a corresponding attenuation of Wnt/β-catenin signalling at day 50, but a reversed signature at day 90 (Supplementary Fig. 3). By sharp contrast, this standard scenario was completely altered in *Eng* haploinsufficient animals. After clipping stimulation, *Eng*^+/−^ littermates showed continuous hair growth during the refractory telogen, despite being defined by high Bmp4 expression, but a significant delayed response during the competent telogen (Fig. 3B), marked by lower Bmp4 expression, a situation mirrored by an altered molecular background signature of the effectors in each time point (Supplementary Fig. 3). These results are in agreement with a defective transmission of the Bmp/Smad signal in *Eng*^+/−^ animals.

**Figure 3.**
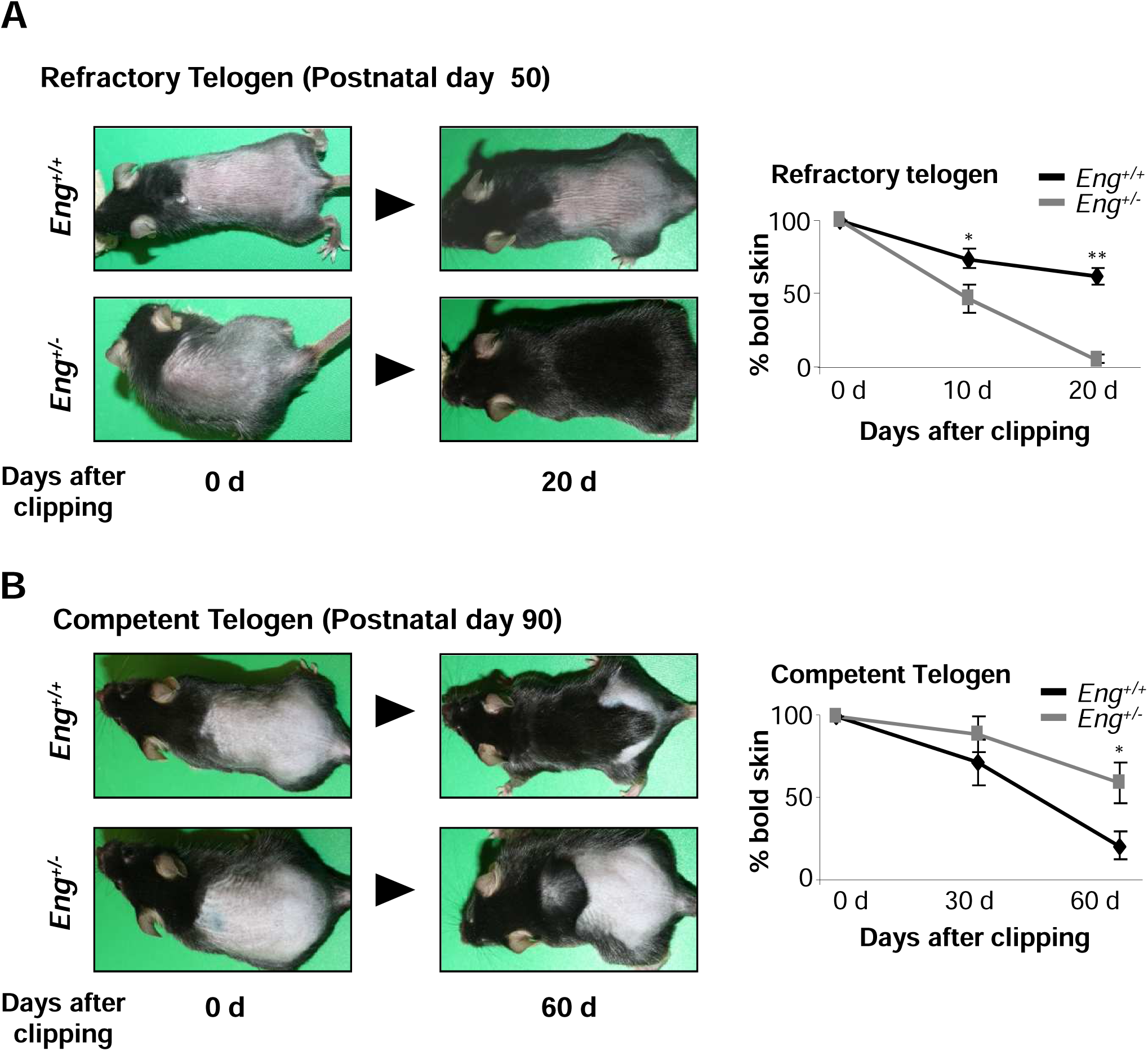
Hair growth is strongly accelerated during the refractory telogen, but is significantly delayed during the competent telogen in *Eng* haploinsufficient mice after clipping stimulation. Induction of hair growth after dorsal hair clipping in *Eng*^+/+^ and *Eng*^+/−^ mice during **A**) the anagen/refractory telogen transition (postnatal day 50), or **B)** the competent telogen phase (postnatal day 90). The experiment finished when most animals of one genotype showed fully completed hair re-growth in the clipped area. Images are representative of three independent experiments including at least three mice for each genotype. The mean percentage of dorsal skin bold area during the experiment +/− SEM is represented; *, p<0.01, **, p<0.005. **C)** Schematic representation of the hair follicle growth cycle and the main signaling pathways involved. Anagen stimulating signal (Wnt/β-catenin) is represented in red, and telogen Bmp/Smad inhibitory signal is represented in green.

To further refine this striking observation, we performed a series of large scale analysis of gene expression using mRNA microarrays and back skin target samples obtained at 50 and 90 days after birth in basal conditions. Interestingly, we found that the large scale gene expression signature was essentially equivalent in *Eng*^+/+^ and *Eng*^+/−^ littermates at the entry of the refractory telogen (postnatal day 50) but differed significantly in the competent telogen/anagen transition (postnatal day 90), when hair follicle stem cell niches are able to respond to stimulatory signals (Plikus et al., 2008) (Supplementary Fig. 4A). At postnatal day 90, several genes, belonging to different gene ontology groups, were found differentially up or down regulated in either *Eng*^+/+^ or *Eng*^+/−^ animals, many of them related to the regulation of hair follicle growth (Supplementary Fig. 4B, C). These results are in close agreement with the *Eng* expression profile during the hair follicle cycle, showing a low expression level at the entry into refractory telogen (postnatal day 50) and a high expression in the competent telogen/anagen transition (postnatal day 90). In addition, these results fit well with the observation that *Eng*^+/+^ and *Eng*^+/−^ resting hair follicles contain a similar number of resident stem cells that deficiently respond to a proliferative stimulus in Eng haploinsufficiency conditions. These observations suggest that a right Eng expression level is required at specific time points of the hair follicle to establish an adequate gene expression pattern.

### Eng haploinsufficiency is associated with a defective proliferative response to growth stimulation of hair follicle bulge stem cells

Taking into account that the hair follicle growth cycle is ultimately regulated by the activity of skin stem cells, we next investigated the effect of a functional reduction of Eng in the proliferative potential of the bulge skin stem cell population. To this end, we first proceeded to identify bulge stem cells as label retaining cells (LRCs) in pulse and long-chase experiments with the nucleotide analogue 5-bromo-2′-deoxyuridine (BrdU) (Braun et al., 2003). We quantified the number of LRCs in the bulge region of hair follicles at 55 days after birth, during which time hair follicles are in the telogen or resting phase and are insensitive to stimulatory signals (Mu et al., 2001). Under these resting conditions, the number and location of LRCs in the bulge region of hair follicles were essentially equivalent in *Eng*^+/+^ and *Eng*^+/−^ littermates (Fig. 4A). Next, we used the phorbol ester TPA to stimulate cell proliferation in the skin and in the hair follicle (Braun et al., 2003; Cotsarelis et al., 1990; Espada et al., 2008). Upon stimulation, wild-type animals showed a significant increase of LRCs in the bulge region, as expected (Fig. 4A). However, *Eng*^+/−^ animals significantly failed to trigger LRC proliferation in the hair follicle (Fig. 4A). These results were corroborated by using a different procedure to activate bulge stem cell proliferation, namely the induction of a transient reactive oxygen species (ROS) production in the skin (Carrasco et al., 2015) (Supplementary Fig. 5), suggesting that *Eng* haploinsufficiency directly affects the proliferative response of skin stem cells to growth signals.

**Figure 4.**
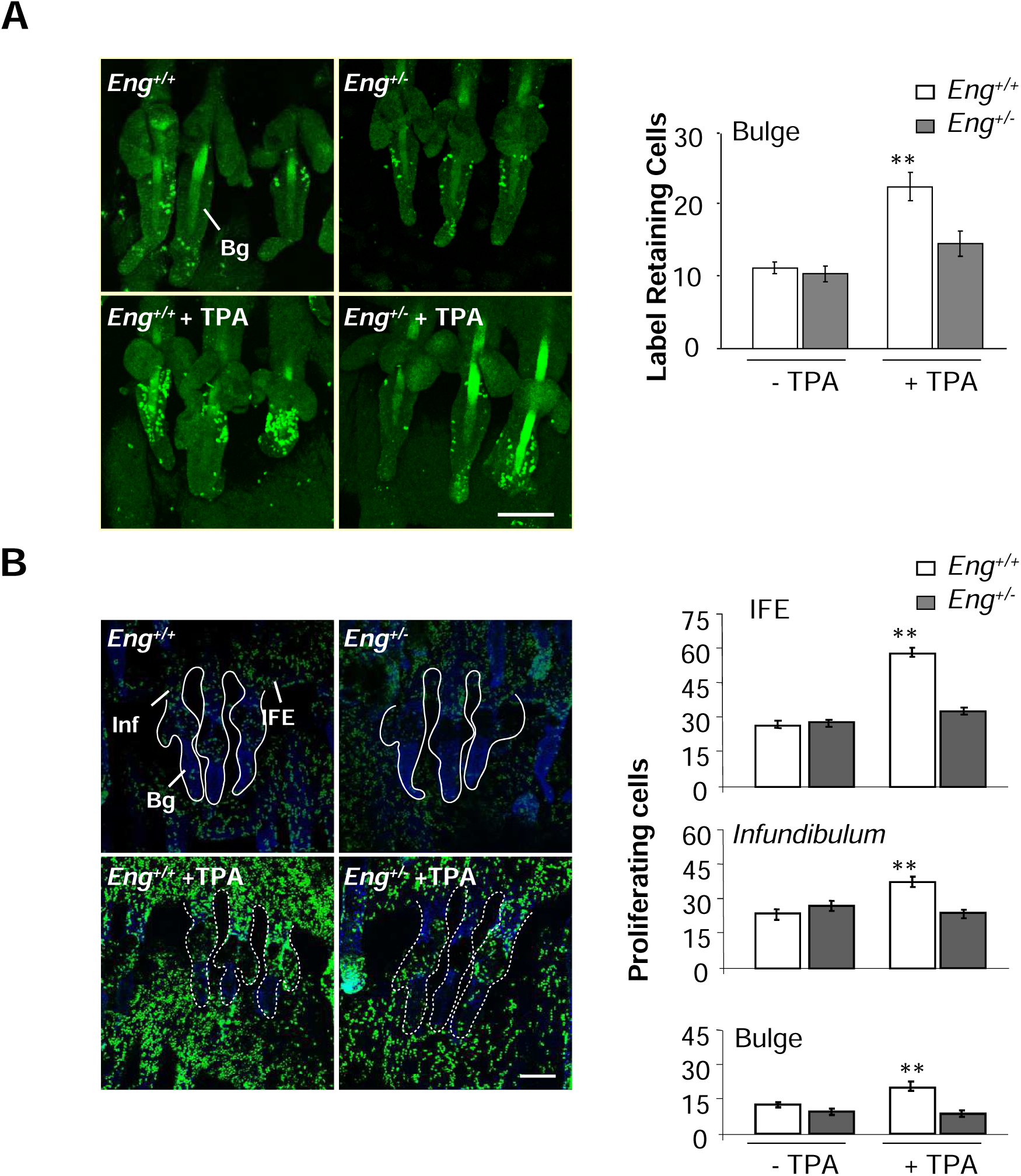
Endoglin haploinsufficiency is associated with a defective proliferative response in the mouse skin and hair follicle bulge stem cell niche after cell growth stimulation. **A)** Quiescent bulge (Bg) stem cells in resting tail hair follicles (telogen, postnatal day 50) were identified as BrdU label retaining cells (LRCs) after a long chase period following neonatal labelling (left panels) and quantified (right panels; the mean +/− SEM is represented, n=3; **, p<0.005). In basal conditions, *Eng*^+/+^ and *Eng*^+/−^ mice showed a similar number of LRCs in the bulge region. However, the proliferative response of these cells after cell growth stimulation with the phorbol ester TPA, identified as a significant increase in the number of LRCs, was severely impaired in *Eng*^+/−^ mice. **B)** Proliferating cells in resting tail skin and hair follicles were identified as positive BrdU cells after a short labelling pulse in adult animals (left panels) and quantified (right panels; the mean +/− SEM is represented, n=3; **, p<0.005). A widespread defective proliferative response after cell growth stimulation with TPA was observed in *Eng*^+/−^ animals in most skin locations, including the interfollicular epithelium (IFE), the infundibulum (Infd) and the bulge region (Bg). In all cases representative confocal microscopy images (maximum projections) corresponding to whole-mounts of tail skin epidermis are shown. Bars: 100μm.

To further confirm the loss of proliferative potential in the skin and, particularly, in the bulge region of the hair follicle in *Eng*^+/−^ mice, we quantified the number of proliferating cells after a short BrdU pulse in adult animals before and after TPA stimulation in different regions of telogen hair follicles and in the interfollicular epidermis (IFE). We found that in the absence of a proliferative stimulus, the number and location of proliferating cells were similar in *Eng*^+/+^ and *Eng*^+/−^ mouse epidermis (Fig. 4B). However, after TPA stimulation, a significantly reduced number of proliferating cells in the bulge region, the infundibulum, and also in the IFE was recorded in *Eng*^+/−^ mice, as compared to control *Eng*^+/+^ littermates (Fig. 4B). We also performed TUNEL assays to rule out the possibility of cell death induction as cause for the loss of proliferative potential in the skin of *Eng*^+/−^ mice (Supplementary Fig. 6). As a whole, these results indicate that Eng haploinsufficiency does not alter the number of resident hair follicle bulge stem cells in a resting (telogen) phase but is associated to a defective proliferative response after cell growth stimulation, e. g., after the induction of anagen entry.

### Eng haploinsufficiency reveals a cross-talk mechanism between Wnt/β-catenin and Bmp/Smad signalling pathways during the hair follicle cycle

In our biological model, *Eng* gene expression is activated about postnatal day 50 (autonomous anagen/refractory telogen transition) (Fig. 1A; Supplementary Fig. 1) after Wnt/β-catenin signalling was operational and Bmp/Smad signalling has been recently activated to slow down hair growth (Plikus et al., 2008). From this point, *Eng* expression steadily increases throughout the telogen phase and drastically decreases at the competent telogen/propagant anagen transition, after that point, in propagant anagen, Wnt/β-catenin signalling is re-activated (Plikus et al., 2008) while Bmp/Smad signalling is completely switched off (Fig. 1A; Supplementary Fig. 1). Thus, the Eng promoter is transcriptionally active when both Wnt/β-catenin and Bmp/Smad signals are switched on; this transcriptional activity is maintained when Wnt/β-catenin signal is switched off, continued strongly repressed when Bmp/Smad signalling is switched off, and increases again its expression when Wnt/β-catenin signalling is reactivated (Fig. 1A).

In this context, we hypothesized that an Eng-dependent cross-talk mechanism between Wnt/β-catenin and Bmp/Smad signalling may act at this point of the hair follicle cycle, a function predicted for feedback target regulators in the dual oscillator model of hair follicle dynamics (Baik et al., 2016; Plikus et al., 2011; Tasseff et al., 2014). We first wondered if β-catenin could interact with the *Eng* promoter as part of a molecular toggle to interconnect both signalling pathways. Indeed, we found that β-catenin strongly binds the *Eng* promoter transcription start site (−52/+132) in the skin of *Eng*^+/+^ mice during autonomous anagen/refractory telogen transition (postnatal day 50), but not during the competent telogen/propagant anagen transition (postnatal day 90) (Fig. 5A). As expected, this binding was not observed in *Eng*^+/−^ littermates (Fig. 5A). These observations are in close agreement with the pattern of *Eng* expression during the hair follicle, and suggest that Bmp/Smad signalling is required for the binding of β-catenin to the *Eng* promoter. This is confirmed by the fact that in *Eng*^+/−^ animals, a reduction of functional Eng (Fig. 1A) impairs Bmp signalling (Fig. 2A), despite expressing similar levels of Bmp during the hair cycle to their *Eng*^+/+^ littermates, and this coincides with the absence of the interaction of β-catenin to the *Eng* promoter (Fig. 5A).

**Figure 5.**
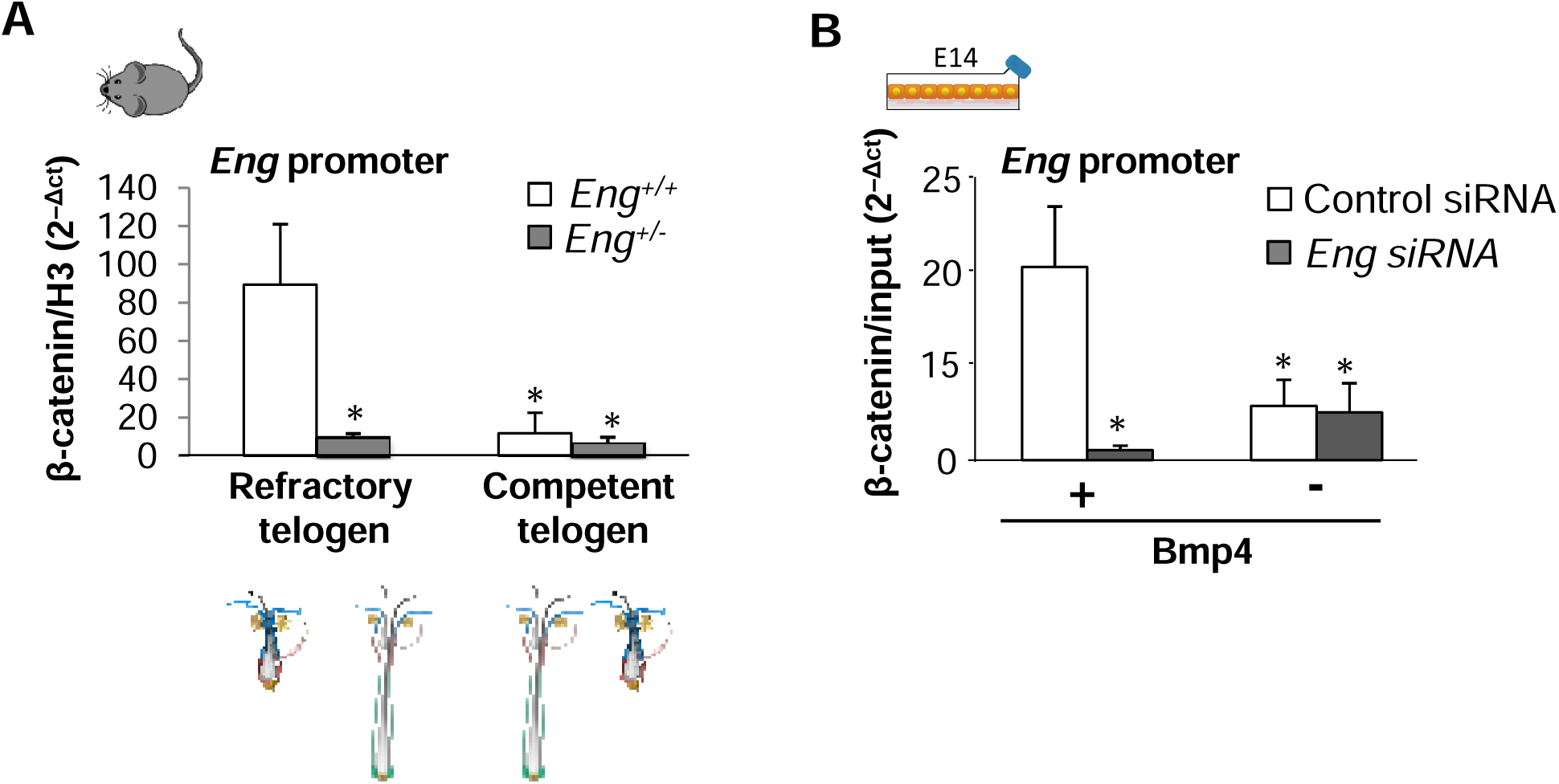
β-catenin directly binds to the Eng promoter depending on Bmp4 signalling and *Eng* expression. Analysis of the interaction of β-catenin with the *Eng* promoter transcriptional start site (−52 / +132 bp) by quantitative ChIP in **A)** *Eng*^+/+^ and *Eng*^+/+^ mice samples of dorsal skin during the anagen/refractory telogen transition (postnatal day 50), or the competent telogen phase (postnatal day 90) and **B)** E14 mES cells transiently transfected with *Eng* siRNA or Control siRNA, and treated with Bmp4 or vehicle control to mimic, respectively, *Eng*^+/−^ and *Eng*^+/+^ tissue microenvironment during refractory or competent telogen. ChIP assays were normalized to histone H3 **(A)** or to input signals **(B)**. Results shown are representative of three experiments in triplicate samples. The mean ± SE is represented; *, p<0.01.

To corroborate the feasibility of this molecular mechanism in a tissue independent context, we used the mouse embryonic E14 cell line. This cell line shows a constitutive accumulation of cytoplasmic β-catenin, driven by LIF-containing media (Takao et al., 2007), mimicking the steady levels of this protein that are observed during the hair follicle cycle in mouse back skin. In this molecular background, we treated E14 cultures with Bmp4 to mimic the anagen/refractory telogen transition and the *Eng* mRNA was depleted by using siRNA interference to mimic Eng haploinsufficiency (Fig. 5B; Supplementary Fig. 7). We found that this model system suitably recapitulated the binding pattern of β-catenin to the *Eng* promoter observed in mouse skin, requiring Bpm4 and Eng expression to occur (Fig. 5B).

The requirement of Bmp4 for the binding of β-catenin to the *Eng* promoter suggests that Smad4, the common transducer of Bmp signalling that is translocated to the nuclear compartment to activate specific gene expression (Blitz and Cho, 2009; Heldin et al., 1997; Lebrin et al., 2004), could be involved in this mechanism. We reasoned that an interaction between β-catenin and Smad4 could drive *Eng* promoter activation. To test this hypothesis, we first performed a series of co-immunoprecipitation experiments in skin samples obtained in the anagen/refractory telogen transition (postnatal day 50) or in the competent telogen/propagant anagen transition (postnatal day 90). The results obtained indicated that in *Eng*^+/+^ mice, β-catenin strongly interacts with Smad4 at the onset of the refractory telogen, coinciding with the co-expression of both proteins in the tissue (Fig. 6A). This specific interaction was lost in *Eng*^+/−^ littermates (Fig. 6A), mirroring the loss of β-catenin binding to the *Eng* promoter. The interaction between β-catenin and Smad4 was also significantly diminished at day 90, during the competent telogen in both *Eng*^+/+^ and *Eng*^+/−^ littermates (Fig. 6A), when Bmp/Smad and Wnt/β-catenin pathways are not activated (Fig. 2A; Supplementary Fig.1). In addition, this pattern of β-catenin/Smad4 interactions in mouse skin was recapitulated in the E14 cell model (Fig. 6B). It is to note that both in mouse skin and E14 cells the interaction β-catenin/Smad4 in co-immunoprecipitation assays was easily observed using an anti-Smad4 antibody but to a lesser extent using an anti-β-catenin antibody. This is an interesting observation since this anti-β-catenin catenin antibody has been previously used in numerous studies to report the binding of β-catenin with different partners (Espada et al., 1999; Espada et al., 2005). As a whole, these results suggest that Eng is essential for both β-catenin promoter binding and β-catenin/Smad4 interaction processes, probably due to its key role as a component of the Bmp receptor complex.

**Figure 6.**
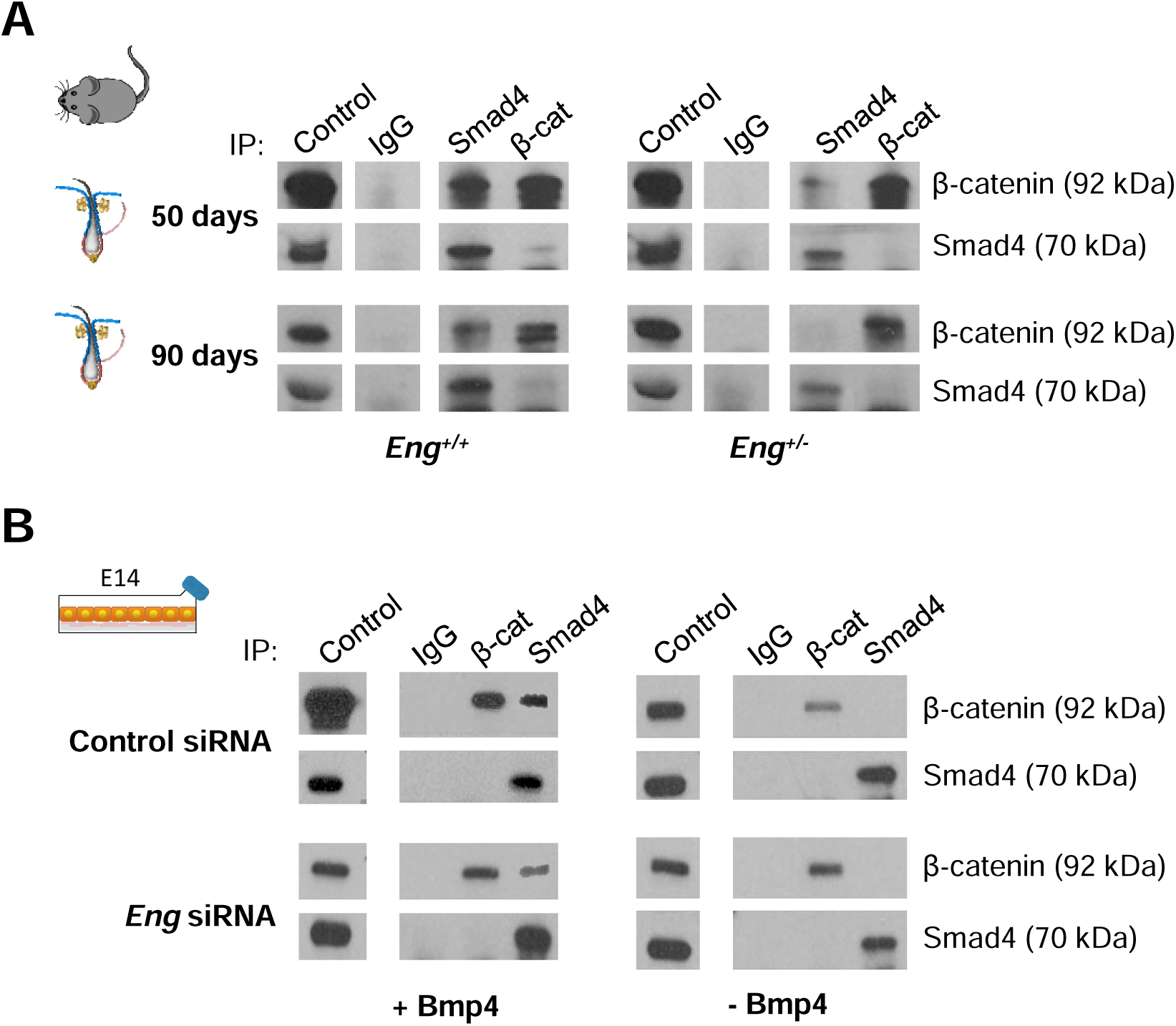
β-catenin interacts with Smad4 depending on Bmp4 signalling and *Eng* expression. Analysis of the interaction between β-catenin and Smad4 proteins by co-immunoprecipitation analysis in **A)** *Eng*^+/+^ and *Eng*^+/−^ mice samples of dorsal skin during the anagen/refractory telogen transition (postnatal day 50), or the competent telogen phase (postnatal day 90) and **B)** E14 cell cultures transiently transfected with *Eng* siRNA or Control siRNA, and treated with Bmp4 or vehicle control to mimic, respectively, *Eng*^+/−^ and *Eng*^+/+^ tissue microenvironment during refractory or competent telogen. Results shown are representative of three experiments, using pools of triplicate samples in each lane.

In this context, we finally tested the potential of β-catenin, Smad4 or a combination of both proteins to regulate the transcriptional activity of the *Eng* promoter. To this end, we performed a series of promoter activation assays in human embryonic kidney 293T cells. This cell line does not express significant levels of endogenous β-catenin, Smad4 or Eng (Fig. 7A), providing a suitable model to analyse the effect of these proteins on the activity of the *Eng* promoter. In this cellular background, expression of activated β-catenin increases Eng protein levels, and this effect is synergistically potentiated by co-expression of Smad4 (Fig. 7A). Moreover, we found that activated β-catenin efficiently activates a reporter plasmid containing the ‒2450/+350 bp promoter region of the *Eng* gene and this activation was strongly increased when both activated β-catenin and Smad4 were co-expressed (Fig. 7B). Interestingly, a high Eng expression is observed in the mouse back skin when high levels of β-catenin and Smad4 are present in the tissue (refractory telogen, postnatal day 70), but a significant reduction of Eng expression is observed when Smad4 levels decay (propagant anagen, postnatal day 100) (Fig. 7C). These results indicate that the β-catenin/Smad4 heterodimer is an efficient activator of the *Eng* promoter transcriptional activity.

**Figure 7.**
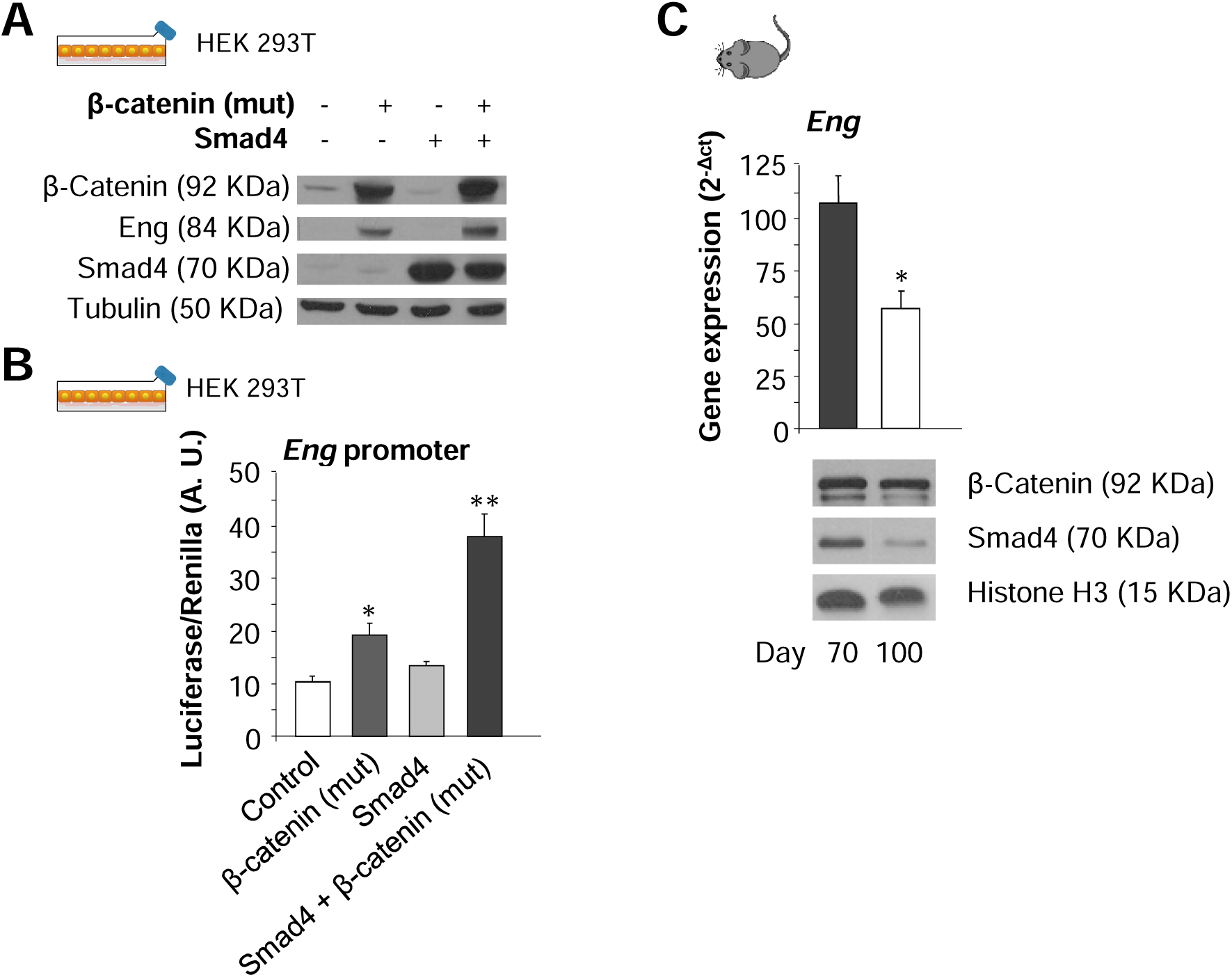
β-catenin and Smad4 synergistically promote the transcriptional activation of the *Eng* promoter and subsequent Eng protein expression. **A)** Immunoblot analysis of β-catenin, Smad4 and Eng protein expression levels in 293T cells after transient transfection with β-catenin and Smad4 expression vectors. Tubulin expression was used as loading control. Results are representative of three experiments, using triplicate samples. **B)** Luciferase reporter assays of *Eng* promoter activity in 293T cells after transient transfection of β-catenin and Smad4 expression vectors in 293T cells. The mean ± SE of Luciferase/Renilla normalized signals (arbitrary units; A. U.) of three independent experiments is represented. **C)** *Eng* mRNA expression quantification by qRT-PCR, normalized to 18S rRNA, in wild-type mouse dorsal skin during the refractory telogen (postnatal day 70) or the propagant anagen (postnatal day 100) of the hair growth cycle (upper panel; the mean +/− SEM is represented, n=3; *, p<0.01) and immunoblot analysis of β-catenin and Smad4 at the same time points, using Histone H3 as loading control (lower panels; results are representative of triplicate samples).

## DISCUSSION

Here we provide evidence of an important role for the Tgf-β/Bmp co-receptor Eng in the molecular switch between Wnt/β-catenin and Bmp/Smad signalling that occurs during the hair follicle growth cycle. We have found that Eng shows a highly defined expression pattern in the skin related to the hair growth cycle phase and to the proliferative state of the resident skin stem cell population, suggesting that Eng levels are an important factor on the whole homeostatic regulation of this tissue. These observations are in agreement with other reports demonstrating that Eng expression can oscillate in the endothelium to act as a modulatory switch between angiogenesis and quiescence programs in the tissue, promoting either Tgf-β/Alk5 or Tgf-β/Alk1 signalling, respectively (Botella et al., 2001; López-Novoa and Bernabeu, 2010; Paus et al., 1999). Mutations in *Eng* causing the autosomal dominant bleeding disorder hereditary haemorrhagic telangiectasia type I (HHT1) are associated to a severe alteration of this signalling balance (Kapur et al., 2013; Lebrin et al., 2004; López-Novoa and Bernabeu, 2010; McAllister et al., 1994; Sanz-Rodriguez et al., 2004). The reduced proliferative potential of bulge LRCs after a growth stimulus is also in close consonance with the finding that reduced Eng expression is associated to a delayed wound healing in the skin (Pérez-Gómez et al., 2014).

Both the hair follicle cycle-dependent expression pattern of Eng in the skin and its involvement in the hair follicle proliferative stem cell response, identifies Eng as a potential feedback regulator target predicted in a coupled dual oscillator model of the hair follicle growth cycle (Tasseff et al., 2014). In this model, cyclic hair follicle dynamics is precisely described and recapitulated as the result of two interacting cell populations, mesenchymal (niche cells) and epithelial (stem cells), showing an out-of-phase synchronized pattern of gene expression. The growth of the hair follicle is mainly driven by the epithelial cell population and this activity is regulated in a feedback pattern by the background mesenchymal cell population through direct cross-talk mechanisms. In this model, it is predicted that the feedback cross-talk between both populations will be mainly regulated by key targets, further identified according to previously published data of whole gene expression patterns in mouse skin. Using this novel approach, Eng was identified as one of these key targets, and the results that we report in this work fully support the role for this protein in the skin predicted by the coupled dual oscillator model.

In this sense, we have accordingly found that a reduced level of *Eng* expression in the skin is associated with the deregulation of the hair follicle cycle, resulting in a delayed establishment of refractory telogen phase. Moreover, we have shown that stimulation of hair growth during the refractory or competent telogen phases of the hair follicle has strikingly different outputs depending on the *Eng* expression background. Interestingly, alterations in the hair cycle associated to mutations in cytokines and receptors involved in Tgf-β/Bmp signalling have been previously reported. Thus *Tgf-β2* mutations result in a delayed anagen establishment (Oshimori and Fuchs, 2012) while *Tgf-β1*^−/−^ mice show a delayed catagen establishment during the first postnatal cycle (Foitzik et al., 2000; Lin and Yang, 2012).Also, over-expression of the Bmp antagonist Noggin in the mouse epidermal basal layer results in a dramatically shortened refractory telogen phase and an accelerated propagation of the hair follicle regenerative wave (Lin and Yang, 2012; Plikus et al., 2008), while subcutaneous Bmp4 injections inhibit anagen establishment (Plikus et al., 2008). These observations point to an essential role for Tgf-β/Bmp in the feedback mechanisms that regulate the inhibition of hair growth, and suggest that Eng, as a part of the Tgf-β/Bmp receptor complex (Cheifetz et al., 1992; Shi and Massagué, 2003), which in turn controls the establishment of the refractory telogen (Plikus et al., 2008), is a central element in the regulation of hair follicle dynamics.

Different reports point to a cross-talk between Wnt/β-catenin and Tgf-β/Bmp signalling as a basic regulatory mechanism of stem cell function in mammalian tissues; for instance, in the regulation of intestine stem cells proliferation (He et al., 2004; Kühl and Kühl, 2013), epidermal/hair placode fate (Fuchs, 2007), or, more recently, in haematopoiesis, where Eng integrates Bmp and Wnt signalling to induce that process (Baik et al., 2016). However, the underlying molecular mechanisms are not well characterized particularly in the hair follicle cycle. Here we have used the theoretical framework of the coupled dual oscillator model of hair follicle dynamics (Tasseff et al., 2014) to identify Eng as a potential key regulator of the feedback cross-talk between these signalling pathways. Supporting this notion, we have found that, in mouse skin, Eng expression increases after anagen establishment, a time point in which β-catenin becomes transcriptionally competent due to Wnt signalling activation (Fuchs, 2007; Kühl and Kühl, 2013). Moreover, our results show that during the mouse hair follicle cycle β-catenin directly interacts with the transcription start site of *Eng* promoter in an Eng/Bmp4 dependent manner, and this binding is associated to *Eng* transcription. These results are fully recapitulated in the E14 embryonic cell model, suggesting that this is a universal mechanism. In addition, we have also found that in both mouse skin and E14 cells, β-catenin interacts with the Bmp signal transducer Smad4 in an Eng/Bmp4 dependent manner, and that both proteins can act synergistically to activate the *Eng* promoter. These observations are in agreement with the fact that *Eng* expression can be regulated in a feedback pattern by the signalling pathways in which it is involved (Baik et al., 2016; Botella et al., 2001).

As a whole, the results reported here indicate that Eng is a key component of the molecular oscillator that regulates the hair follicle cycle, acting as an important element in the dynamic cross-talk between Wnt/β-catenin and Tgf-β/Bmp signalling. This cross-talk would consist in a feedback mechanism in which Eng expression is regulated by β-catenin binding to the *Eng* promoter in a pattern dependent on Bmp4 signalling and further driven by the interaction of β-catenin with the Bmp4 transducer Smad4, and in turn, as recently has been demonstrated, Eng induction enhances Wnt activity, that promotes the stabilization of activated Smads (Baik et al., 2016), establishing a negative feedback that points Eng as a potential universal oscillator. As Eng is required to transmit the Bmp4 signal and to mobilise Smad4, a β-catenin-dependent transcriptional activation of *Eng* expression suitably close the feedback circuit. It is tempting to speculate that a similar or equivalent mechanism could be found in mammalian tissues in which a coordinated Wnt/β-catenin and Tgf-β/Bmp/Smad signalling cross-talk is involved in the regulation of tissue homeostasis, bringing to the forefront a new research framework to investigate adult stem cell biology.

## ACKNOWLEDGEMENTS

This work has been supported by grants from Ministerio de Economía y Competitividad of Spain (RTC-2014-2626-1 to JE, SAF2013-46183-R to MQ and SAF2013-43421-R to CB), Comunidad de Madrid (S2010/BMD-2359, SkinModel to JE and MQ), Instituto de Salud Carlos III (PI15/01458 to JE) and Centro de Investigación Biomédica en Red de Enfermedades Raras (CIBERER; ISCIII-CB06/07/0038 to CB) financed jointly by the European Regional Development Funds (FEDER). EC and MIC were supported by Spanish MECD-FPU and UAM-FPI fellowships, respectively. We thank Michelle Letarte (The Hospital for Sick Children, Toronto, Canada) and Jose Miguel Lopez Novoa (Universidad de Salamanca, Salamanca, Spain) for kindly providing the *Eng* haploinsufficeint strain

## CONFLICT OF INTEREST

The authors declare no competing financial interest.

